# Automated Skin Lesion Detection and Prevalence Estimation in Tamanend’s Bottlenose Dolphins

**DOI:** 10.1101/2024.07.15.603578

**Authors:** Colin J. Murphy, Melissa A. Collier, Ann-Marie Jacoby, Eric M. Patterson, Megan M. Wallen, Janet Mann, Shweta Bansal

## Abstract

Anthropogenic global change is occurring at alarming rates, leading to increased urgency in the ability to monitor wildlife health in real time. Monitoring sentinel marine species, such as bottlenose dolphins, is particularly important due to extensive anthropogenic modifications to their habitats. The most common non-invasive method of monitoring cetacean health is documentation of skin lesions, often associated with poor health or disease, but the current methodology is inefficient and imprecise. Recent advancements in technology, such as machine learning, can provide researchers with more efficient ecological monitoring methods to address health questions at both the population and the individual levels. Our work develops a machine learning model to classify skin lesions on the understudied Tamanend’s bottlenose dolphins (*Tursiops erebennus*) of the Chesapeake Bay, using manual estimates of lesion presence in photographs. We assess the model’s performance and find that our best model performs with a high mean average precision (65.6%-86.8%), and generally increased accuracy with improved photo quality. We also demonstrate the model’s ability to address ecological questions across scales by generating model-based estimates of lesion prevalence and testing the effect of gregariousness on health status. At the population level, our model accurately estimates a prevalence of 72.1% spot and 27.3% fringe ring lesions, with a slight underprediction compared to manual estimates (82.2% and 32.1%). On the other hand, we find that individual-level analyses from the model predictions may be more sensitive to data quality, and thus, some individual scale questions may not be feasible to address if data quality is inconsistent. Manually, we do find that lesion presence in individuals suggests a positive relationship between lesion presence and gregariousness. This work demonstrates that object detection models on photographic data are reasonably successful, highly efficient, and provide initial estimates on the health status of understudied populations of bottlenose dolphins.

## 1 Introduction

Globally, direct and indirect anthropogenic impacts on wildlife are increasing at an alarming rate; the rise of emerging infectious diseases, habitat degradation, environmental pollutants, and human-induced climate change threatens their well-being and survival (Barnosky et al., 2011; Gulland & Hall, 2007; Kelly et al., 2021; Pimm et al., 2014; Simeone et al., 2015). These rapid changes underscore the urgency for monitoring wildlife health. While several technological advances allow researchers to amass the necessary data quickly (Allan et al., 2018), big data processing has become a bottleneck that humans cannot manually manage. However, techniques such as machine learning improve data processing times, and applying them to populations of interest may help improve response time to ecological threats (e.g., disease outbreaks, population declines, etc.) (Kelly et al., 2021; Keshavamurthy et al., 2022). Of particular interest is the impact of environmental change on marine species, which remain critically understudied even though more than two-thirds of their habitat is being altered by anthropogenic disturbances (Díaz et al., 2019). Delphinid species (dolphins) face numerous threats with significant population declines (Gulland & Hall, 2007; Simeone et al., 2015). They are also predators and sentinel species essential for healthy ocean ecosystems (Bossart, 2011; Heithaus et al., 2008; Moore, 2008). Given their importance, our work aims to demonstrate how machine learning can improve the effectiveness and efficiency of health monitoring in vulnerable species using data collected on Tamanend’s bottlenose dolphins (*Tursiops erebennus*) as a case study.

Advances in machine learning techniques for data processing and analysis are revolutionizing ecology with tools that enable processing and analysis of increasingly complex data (Christin et al., 2019; Olden et al., 2008). Current applications include object detection algorithms for terrestrial animals that detect the location and types of wildlife present in a region (Chen et al., 2019; Roy et al., 2022), automation of the description of social behavior in animals by describing body position and tracking gazes (Pereira et al., 2019; Turesson et al., 2016), and detecting activity states such as foraging (Browning et al., 2018), among others. Regarding wildlife health, machine learning is used to parse disease reports to track outbreaks as they occur (Kelly et al., 2021) and predict disease outbreaks temporally and spatially (Keshavamurthy et al., 2022).

Machine learning has also begun transforming data processing tasks in the marine environment. It is currently used in passive acoustic monitoring to classify click patterns for population location and behavior (Caruso et al., 2020; Roch et al., 2021), identify dolphins from photographs of dorsal fins (Maglietta et al., 2022), and in transcriptome analysis to investigate the effects of toxins in marine mammals (Mancia et al., 2014). However, despite their vulnerability to global change, researchers have yet to apply these techniques to the ecological and health monitoring of key marine species, such as bottlenose dolphins.

Populations of bottlenose dolphin species, like the Tamanend’s bottlenose dolphin, are found in coastal waters worldwide, making them particularly vulnerable to anthropogenic stressors such as chemical pollution, biotoxins, run-off, noise, and other local impacts (Alter et al., 2010; Page-Karjian et al., 2020; Pierce et al., 2008; M. Van Bressem et al., 2009; Yang et al., 2021), which have been shown to worsen their overall health (Alter et al., 2010; Collier et al., 2022; Deming et al., 2020; Fazioli & Mintzer, 2020; Sousa et al., 2019; Toms et al., 2021). They are also particularly vulnerable to infectious diseases that may cause mass mortalities, such as the 1987 and 2013 dolphin morbillivirus epizootics on the U.S. Atlantic coast, which depleted some populations of dolphins by 40-50% (Lipscomb et al., 1994; Waring et al., 2016).

To effectively monitor disease and health risks for bottlenose dolphins, researchers use several methods. Capture and release assessments currently yield the most detailed picture of health status, as researchers can perform various tests such as blood sampling and surgical biopsy (Barratclough et al., 2019; Kucklick et al., 2011). Remote biopsying of dolphin skin and blubber from boats yields health-related information on pollutants, hormones, and aging (Barratclough et al., 2019). However, these methods are invasive in nature, resource-intensive, require extensive training and permitting, and incur high costs which generally result in a small sample size of individuals tested. These methods have been shown to increase the release of stress hormones in the study organisms, which may alter or skew test results and diminish the accuracy of the assessment (Fair et al., 2014, 2017).

As an alternative, photographic data collection is a more broadly applicable method to monitor the health of cetacean populations. Since bottlenose dolphins frequently surface to breathe, researchers can photograph visual evidence of skin lesions, which are regions of damage or discoloration. Photographs can either be taken with a handheld camera from a boat or with unmanned aerial surveillance (Castrillon & Nash, 2020). Differences between lesions caused by disease or environmental stress can be accounted for by classifying differently shaped and colored lesions, forming a rough picture of etiology (Toms et al., 2020). The cost and invasiveness of photographic data collection are minimal compared to other methods, providing more data over a shorter period of time. However, the data processing of thousands of photographs for skin lesions is labor intensive and subject to error and bias. Some lesions are small, and photo quality can be poor due to focus, glare, water droplets, and exposure.

Furthermore, researcher experience in identifying and classifying lesions might vary, and coupled with subjective bias and different classification criteria this may lead to incongruous data and flawed analysis within study sites and between study sites (Toms et al., 2020). Object detection models, which are trained to detect and classify regions of an image containing objects of interest, may offer an apt solution to these problems by automating data processing and reducing error and bias for better real-time monitoring of bottlenose dolphin health. Of interest would be models designed to detect smaller-scale lesions as they 1) serve as early warning signs of disease such as poxvirus (Geraci et al., 1979; Maldini et al., 2010; Powell et al., 2018; M. F. Van Bressem et al., 1999), and 2) are the hardest to reliably detect and classify given that they often occupy less than 0.01% of a photograph. Detecting lesions efficiently with these models would allow scientists to estimate lesion prevalence in a population and test hypotheses at an individual level, such as the relationship between health and behavior. For example, lesion pathologies might be more prevalent in individuals who have higher rates of contact with conspecifics. Alternatively, outward signs of illness or disease have been known to elicit avoidance behaviors in several species (Stockmaier et al., 2023), meaning that dolphins with lesions may have lower rates of contact.

In this study, we train an object detection model that detects and classifies evidence of small skin lesions present in photographs of Tamanend’s bottlenose dolphins from the Potomac-Chesapeake Dolphin Project (PCDP). We then apply this model to additional PCDP data not used in the training process to test its ability to answer ecological questions at both the population and individual levels. The health of the dolphins in this region is of interest to marine mammal researchers and managers alike; during a 2013 disease outbreak of Mid-Atlantic bottlenose dolphins, the plurality of disease-related deaths (>400) occurred near the PCDP’s main study area (Waring et al., 2016). While lesion presence has been anecdotally observed in the Potomac-Chesapeake from the more than 100,000 photographs collected since 2015 by the PCDP, true lesion prevalence is difficult to estimate. This study, therefore, provides a comprehensive look at the health of the animals that utilize the Chesapeake Bay watershed and improves the efficiency of ecological monitoring in cetaceans more broadly.

## 2 Methods

### 2.1 Dataset

We used data from photographs of bottlenose dolphins from field surveys conducted between 2015 and 2018 with the PCDP.

#### 2.1.1 Data Collection

Surveys were conducted in Maryland and Virginia waters from the lower tidal Potomac River (southeast of the Governor Harry W. Nice Memorial Bridge, N 38.3616, W 76.9971) to the middle Chesapeake Bay (specifically Ingram Bay N 37.8007, W 76.2885). Researchers conducted line transects and searched for dolphins within the established study area; when dolphins were sighted, a survey commenced, where observers took photographs of all present individuals for identification and recorded their predominant behavioral activities (Karniski et al., 2015). We examined approximately 8,000 photographs from 56 surveys conducted between 2015 and 2018 for the training and testing of all models (see Tables S1 and S2 in the Supplement). Each photo received a quality score from zero (lowest quality) to three (highest quality) (see Table S3 in the Supplement) (Urian et al., 2015). Images with a higher photo quality rating were less likely to have lesions obscured by issues such as lack of focus, droplets, shadows, and glare. Only photographs rated two or three in quality were used to train models. Additionally, for making prevalence estimates with our models, we used only images with a single dolphin present as the model was not trained to match lesions to specific dolphin identity within a photograph.

#### 2.1.2 Permits and Ethics Statement

All data was collected passively with no direct animal interactions under NMFS Permits 19403 and 23782. There was no evidence that the presence of our research vessel affected dolphin behavior; to reduce potential stress we took care to approach groups slowly (under 7 kph), kept approximately 10-20m away from a group, and shut off our engine completely when groups stayed in the same area.

### 2.2 Lesion Classification

For this project, we focused on smaller circular lesions. Based on lesion classifications in bottlenose dolphins from the literature (Powell et al., 2018; Toms et al., 2020), we first classified lesion shape as a round spot or a concentric fringe ring. Fringe ring lesions in particular are closely linked with underlying cetacean poxvirus, which also presents as tattoo skin disease (Geraci et al., 1979; Maldini et al., 2010; M. F. Van Bressem et al., 1999). We then classified the lesion’s coloration as dark (hyperpigmented compared to the majority of the dolphin’s coloration) or pale (hypopigmented compared to the majority of the dolphin’s coloration). With fringe rings, we classified the lesion’s color based on the outermost ring. Therefore, we classified four lesion categories: pale spot, dark spot, pale fringe, and dark fringe (Figure 1). Our dataset also contained dolphins with both dark and pale amorphous lesions (i.e., lesions with irregular shapes and sizes). However, we did not include these lesions in our analysis due to their extreme variability and underrepresentation in the dataset (less than 8% of all lesions). Linear lesions such as rake marks or entanglement scars were also not included.

**Figure 1.**
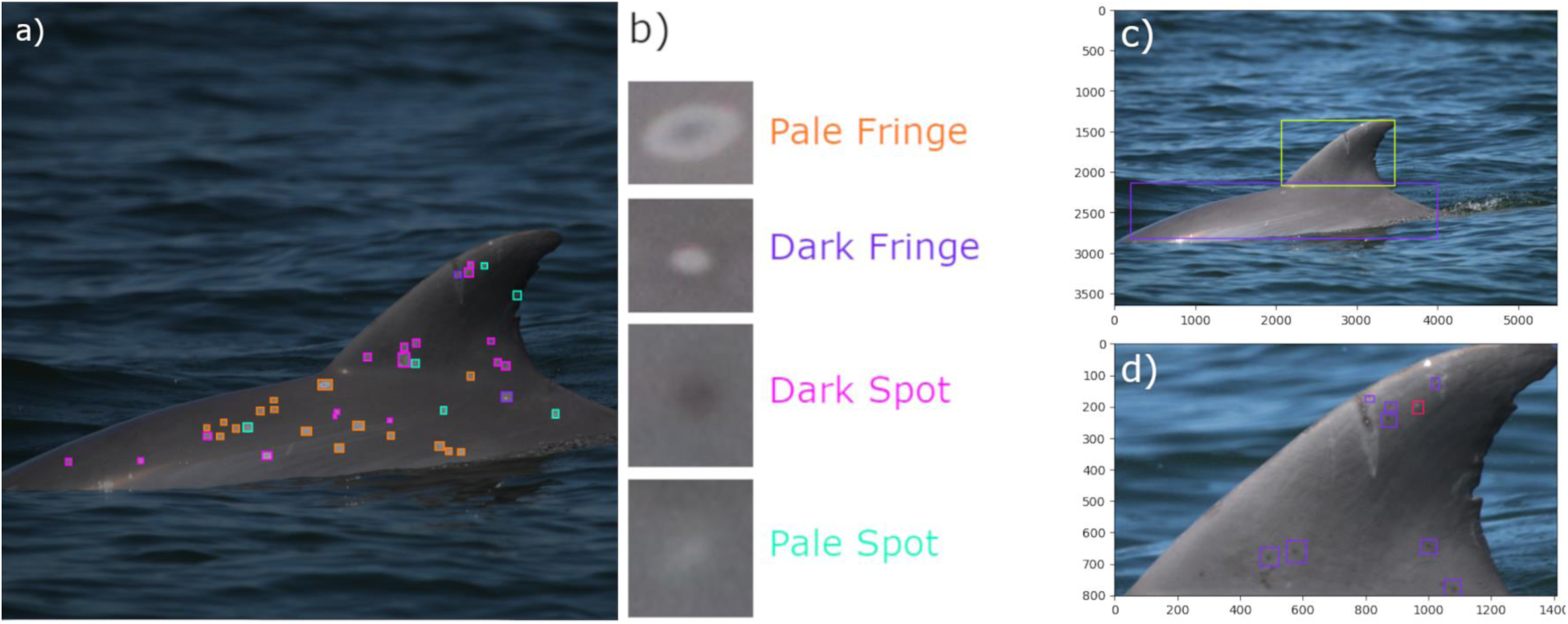
Examples of lesion annotations and detections. Lesion locations were annotated on the Roboflow platform (a) and sorted into four classes (b). Annotations were stored as bounding box coordinates containing the region of interest (a). Also included are example bounding box detections from a detector aimed at locating the dolphin’s body and dorsal fin in the image (c), and those from a detector that identifies spot lesions on a dolphin’s dorsal fin (d).

For each photo in our dataset, two different researchers (one “novice” with under one year of experience, and one “expert” with six years of experience) searched for lesions on dolphin skin. The novice established the initial presence or absence of a skin lesion on a dolphin in each photo, and the expert then confirmed these determinations and further categorized the lesions into the four categories (Figure 1b). In the case of a disagreement in presence/absence of a lesion between the two researchers, we deferred to the call of the expert. We then selected photographs that gave us a balanced set of annotations across all four lesion classifications, yielding 405 images.

For each photograph, we drew bounding boxes around every visible lesion from one of the four lesion categories. The photographs were then split into training data (70%), validation data (20%), and testing data (10%). The model used training data to learn the lesion types, used validation data to prevent overfitting to the training data, and used testing data to provide an assessment of model performance.

### 2.3 Lesion Detection Model

To carry out dolphin lesion detection using our curated dataset, we trained and deployed a computer vision model using the Ultralytics YOLOv5 object detection model within the Roboflow software platform (Jocher et al., 2022). We tested two model frameworks to evaluate small lesion detection using photographic data from the field. The first framework using a single model required more input from the user. Based on the results of the models from the first framework (“one step models”), we then developed a fully automated model framework that leveraged what we learned from the first model (“two step models”). We trained 10 model replicates for each model framework to capture variation.

To assess the performance of our models, we used the mean average precision (mAP), which is derived by averaging the area under the precision-recall curve for each lesion class present in the training data.

#### 2.3.1 Developing a one-step computer vision model for small lesion detection

We first developed models to detect small lesions in a dolphin photograph which we refer to as “one-step models”.

1. *Uncropped model with tiling and augmentation:* Photographs from the field were used in this training. To improve model accuracy, we used tiling, which divides each image into smaller tiles. Additionally, we used random crop, random rotation, random shear, random blur, random noise, and mosaic as image augmentations to improve the ability of our model to generalize and thus perform more effectively on images outside of the training set. Details of these augmentations can be found in the Supplement, with an example of tiling in Figure S1.
2. *Cropped model with augmentation:* All images were cropped before model training to include the whole dolphin with a fixed 1:1 square aspect ratio, aiming to preserve the dolphin’s body in the image while reducing the region of the image not containing a dolphin (water, sky, background, etc.). Tiling was not carried out due to the varying image sizes. The same image augmentation approaches were used as our first model. From this stage onwards, transfer learning was also applied using a specialized prototype model; see Supplement for more details.
3. *Cropped model with partitioning and augmentation:* In our training data, spot lesions are overrepresented compared to fringe rings, which causes lower performance on the classes with lower representation. To correct this issue, we partitioned the training data for spot lesions and fringe ring lesions, and used the cropped model with augmentation as above.

#### 2.3.2 Developing a two-step computer vision model for lesion detection with automated cropping

Our efforts with the one-step model clarified the need for cropping photographs to remove negative space before carrying out small object detection. However, requiring manual cropping of the images meant only part of the analysis process was made more efficient by our models. Thus, we automated the process of cropping by developing another object detection model that locates a dolphin’s dorsal fin and the remaining body in an image and crops the image to those detections. We used 380 photographs annotated with the dolphin’s dorsal fin and the rest of the dolphin’s visible body to train an object detection model to recognize dolphin bodies and dorsal fins (Figure 1c). This body and dorsal fin detection model was trained 10 times to account for potential variation in model performance, and the resulting performance of this model can be found in Table S4.

The body and dorsal fin annotations were then cropped as separate images and used to train new cropped lesion detection models with partitioning and augmentation (as described above in Section 2.3.1 #3). We refer to these as “two-step models.”

### 2.4 Testing the effects of image quality on model performance

We tested how the quality of the photographic data affects lesion detection model accuracy using two aspects of data quality: a) the number of photographs provided per dolphin and b) the quality of the photographs (as defined in Section 2.1). We evaluated the effect of these data quality measures on the performance of the two-step model framework. For this evaluation, we used a dataset of 366 images of 220 individual dolphins that had not been used in the training process with varying numbers of images per dolphin (usually one to three) and varying image quality (2- star and 3-star). We ground truthed the model against a manual lesion determination by two researchers (see Section 2.2) in the 366 images and used a bootstrapping approach to generate a 95% confidence interval on these estimates. The model’s accuracy for both data quality measures was calculated as the proportion of matches between manual and model lesion determinations, i.e., when both the model and the manual determinations are positive or negative for a lesion type. For image quality, these matches were determined by image. For the number of photos per dolphin, these matches were determined by dolphin. To decrease the chance of false positives in this analysis, if there were three or more photos of a dolphin, we ensured that more than 25% of their photos were marked positive for a lesion by the model for a dolphin to be considered positive for that lesion according to the model.

### 2.5 Addressing epizoological questions with the lesion detection model

To test the model’s ability to answer questions of interest at the population level, we used our best lesion detection model to estimate *lesion prevalence* (defined as the number of unique individuals with lesions divided by the total number of observed animals). For this, we used the same set of 366 images of 220 dolphins with manual lesion detections that we used for our tests in Section 2.4.

To test the model’s applicability at the individual scale, we addressed whether the presence of a lesion is associated with the gregariousness of an individual. We defined the average number of individuals each dolphin was sighted with in a survey as a proxy for gregariousness, where individuals with higher values are considered more gregarious. We hypothesized that because fringe ring lesions are thought to be associated with infectious disease, the probability of having a fringe ring lesion would be higher in more gregarious individuals (Powell et al., 2020).

Alternatively, lesions of either type may lead to avoidance behaviors by conspecifics (Stockmaier et al., 2023) or sickness behaviors by the individual, such as reduction in activity and social interactions (Lopes et al., 2021), meaning that less gregarious individuals might be more likely to have either a spot or fringe ring lesion. We ran two generalized linear mixed models with binary response variables for the presence or absence of a lesion on an individual based on 1) our manual determination of either fringe ring or spot lesion presence and 2) our model’s determination of fringe ring or spot lesion presence. We used a continuous predictor variable for our gregariousness proxy, and random effects for the year of the photograph and the identity of the dolphin to control for variability in data collected across years and individual behavior of the dolphin, respectively. We then compared the effect of gregariousness on lesion presence between the manual and the model.

## 3 Results

### 3.1 Dolphin lesion detection is effective with high object coverage

The mean mAP for the uncropped one-step model is 8.5%. Cropping of the photographs to the dolphin significantly improves the performance of the one-step models (mean mAP = 50.5%). Further partitioning of the cropped model into spot-only and fringe ring-only detection further improves the model for fringe ring detection (mean mAP = 72.8%), but not for spot detection (mean mAP = 39.3%) (Figure 2A). This confirms that a high object-to-background ratio is key to improved small object detection for this application.The two-step process (with automatically cropped images to the dolphin body and dorsal fin) greatly improves the performance of the spot detection model on dolphin bodies (mean mAP = 73.1%) and dorsal fins (mean mAP = 69.5%). The fringe ring detection model also significantly improves for both the body (mean mAP = 81.9%) and dorsal fin (mean mAP = 78.9%) (Figure 2B). Results of an ANOVA and Tukey HSD test can be found in Table S5.

**Figure 2.**
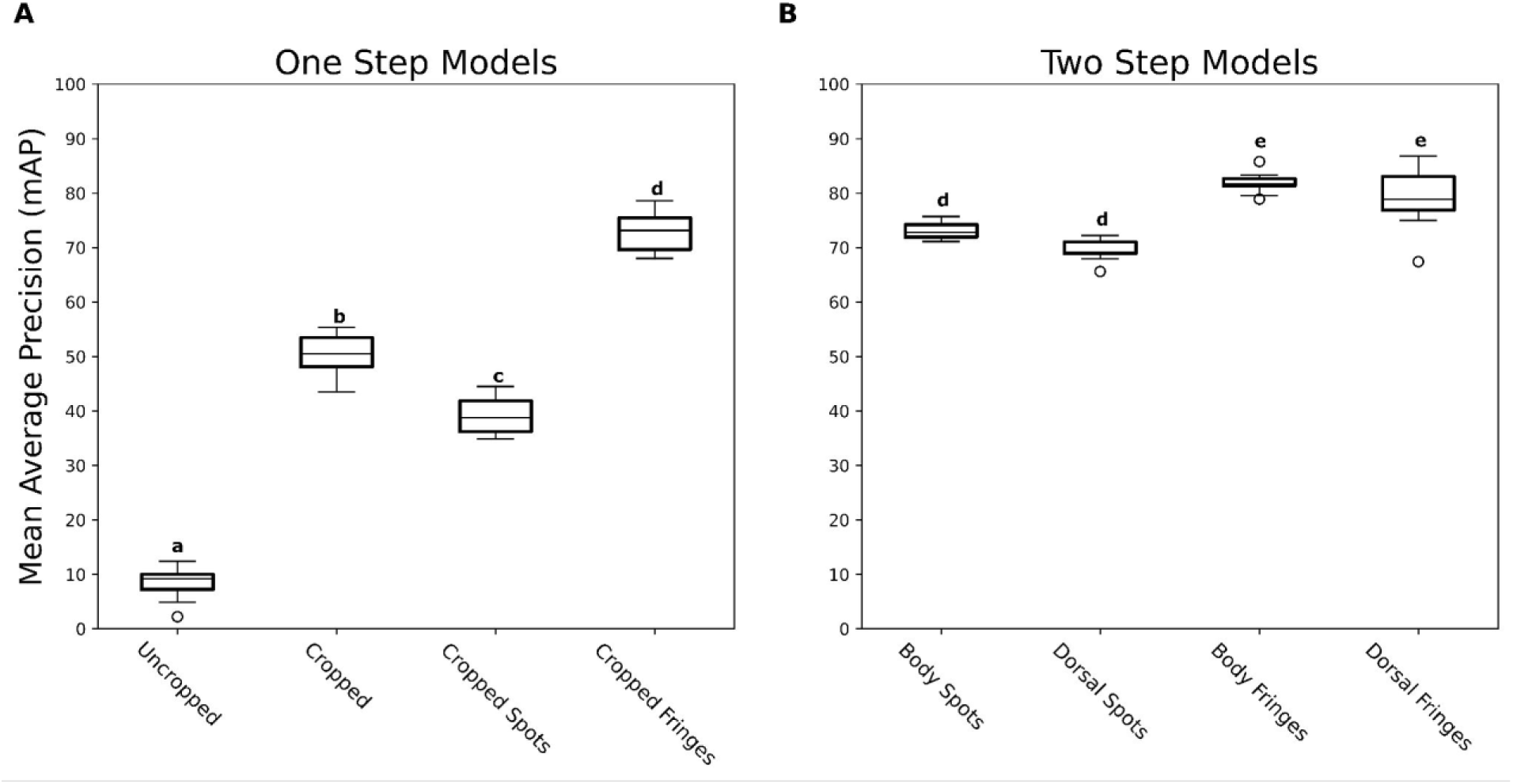
Model training results show higher performance of two-step models. Mean average precision as a measure of model performance for the one-step (Fig 2A) and two-step (Fig 2B) training process. Cropping and partitioning improve the mean average precision (mAP) and two-step models consistently have higher mAP across both lesion types and body parts.

### 3.2 Lesion detection performance varies by data quality

We test how the performance of our model changed with image replication (number of images available per dolphin) and image quality. Accuracy for detecting a spot lesion on a single dolphin significantly increases (69.7% to 85.7%) as the number of photos representing a given dolphin increases (Figure 3A). For fringe ring lesions, we see a smaller increase in accuracy values (75.0% to 78.6%) as the number of photos increases (Figure 3B). For image quality, a statistically significant improvement is seen in the model’s ability to detect lesions in 3-star images (100%) compared to 2- star images for spot lesions (63%) (Figure 3C), but a small decline in performance is seen for the same comparison with fringe ring lesions (2 star: 79.7%, 3 star: 70.6%) (Figure 3D). Significance results are based on a one-way ANOVA and Tukey’s HSD test. A full breakdown of the confusion matrices for all of these tests can be found in Tables S6-9.

**Figure 3.**
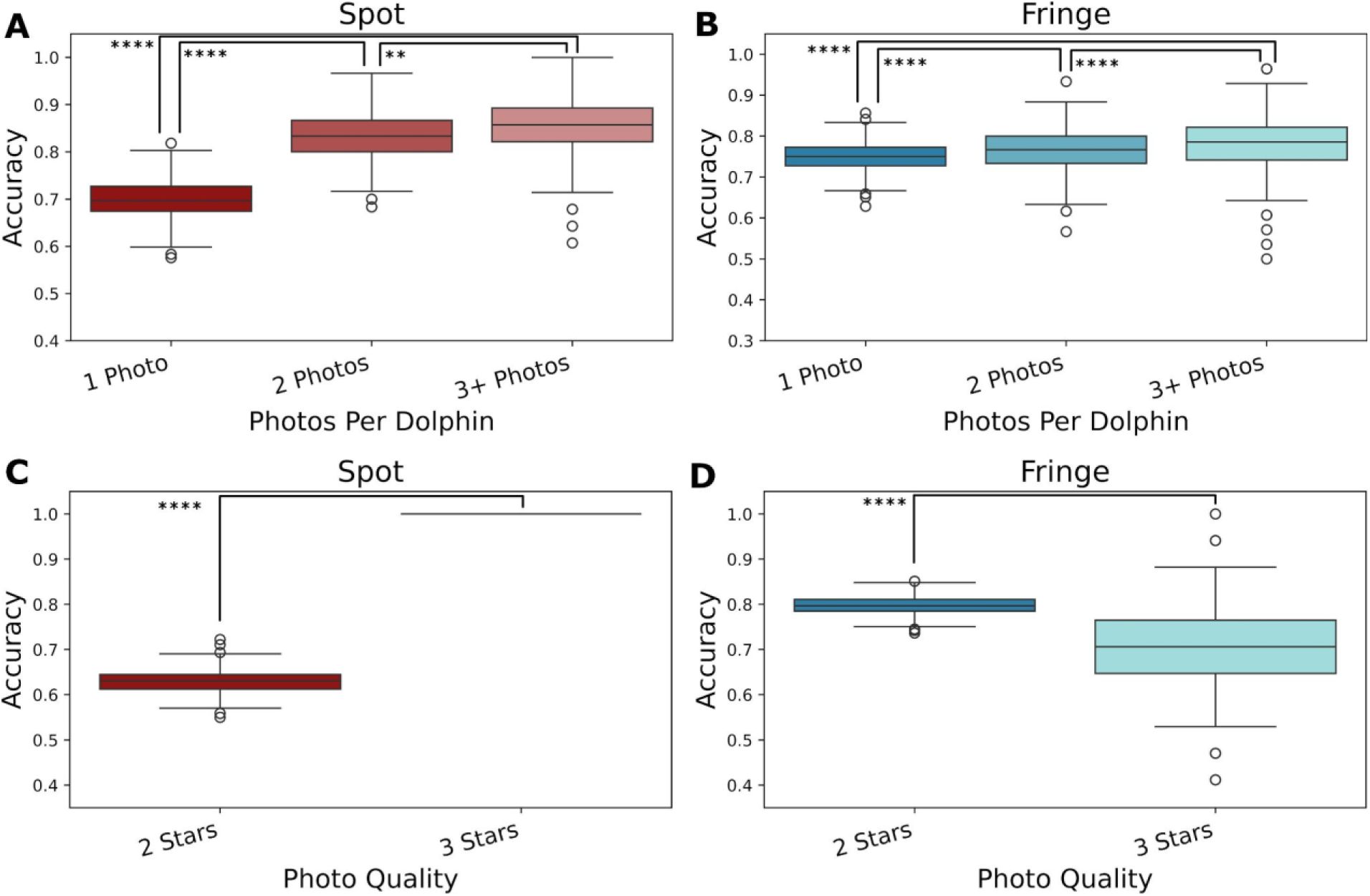
Higher photographic data quality improves performance of spot lesion detection but has unclear effect for fringe ring lesion detection. Our test dataset of images with a single dolphin and verified manual lesion presence was subdivided based on either the number of photos per dolphin for the dolphins represented (A, B) or by the image quality rating of the photos (C, D). Accuracy increases with quality for spot lesions (red: A, C) but the trend is muddied for fringe ring lesions (blue: B, D).

### 3.3 Automated lesion detection reasonably estimates lesion prevalence

Deploying the best model (two-step model) on a set of data with only one dolphin in each photograph (366 photographs and 220 dolphins), we rapidly assessed small lesion prevalence within the sample using the model in 1.5 hours, whereas the manual assessment took over 80 hours between two researchers. The model’s average prediction of prevalence for spots (72.1%) and fringe rings (27.3%) are slight underestimates of the manual predictions (82.2% and 32.1%, respectively) (Figure 4).

**Figure 4.**
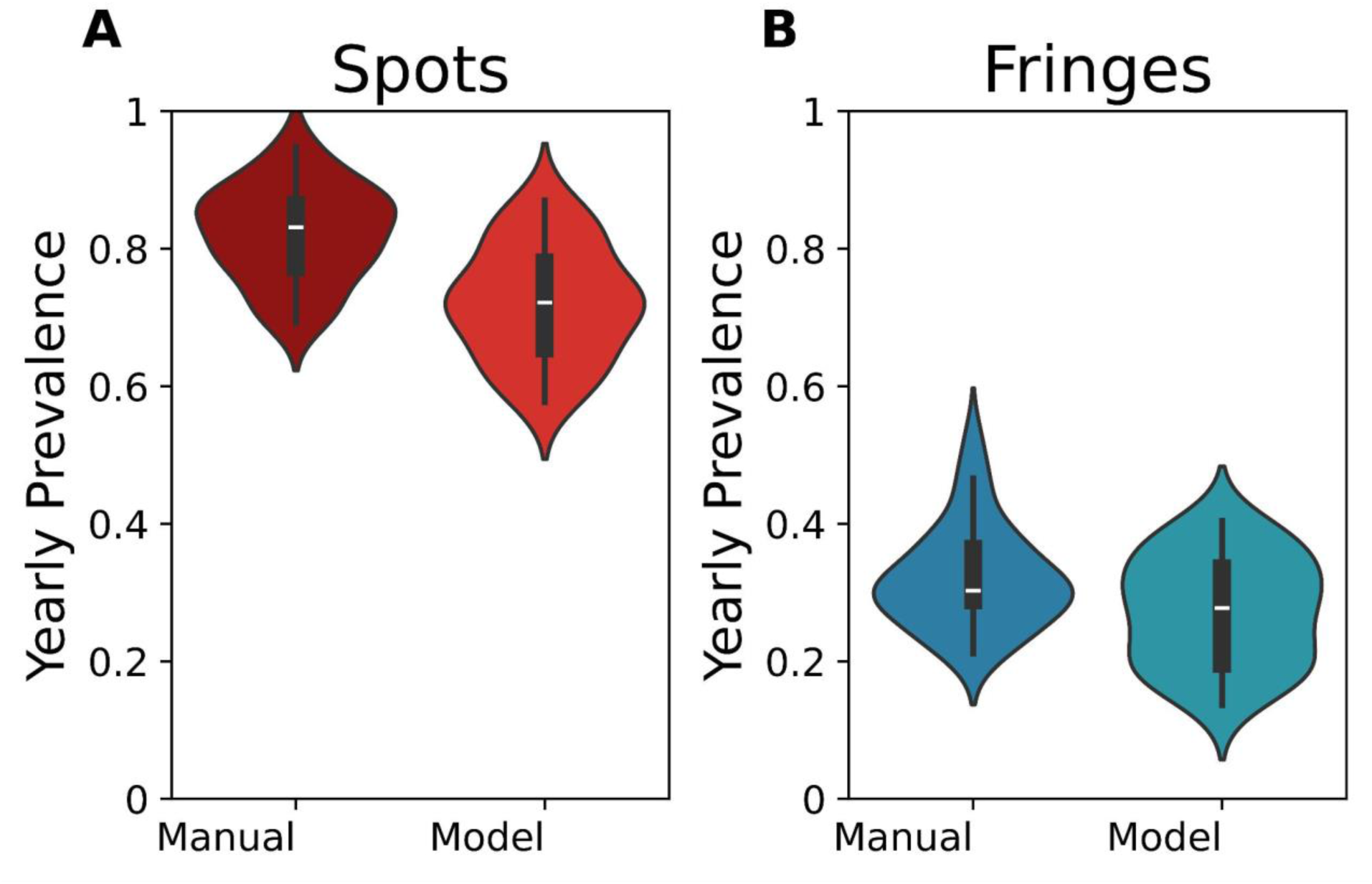
Models slightly underpredict lesion prevalence. The model made predictions for lesion presence by image, summarized into estimates of lesion prevalence for the population by matching predictions to dolphins. The model estimates a slightly lower prevalence than manual estimates of prevalence for the same single-dolphin subset for spots, and is nearly equal for fringe ring lesions.

### 3.4 The ability of automated lesion detection to address individual-scale ecological questions is sensitive to data quality

We seek to characterize the association between individual dolphin gregariousness and lesion presence, as this can inform the role of social behavior in infectious disease risk and the role of adaptive behaviors in the presence of poor health. We find a significant positive effect of gregariousness on the presence of fringe ring lesions when using our manual determination but no effect when using the model’s determination (Figure 5). For spot lesions, there is no significant effect of gregariousness in either the model or manual determination of spot prevalence.We find that the model accuracy of lesion detection (defined as an agreement between the manual and model lesion presence) is lower at larger gregariousness values, which likely drives this difference (see Figure S2). We suggest this is due to poorer data quality at larger gregariousness values, as we see a negative effect between the number of photos per dolphin and gregariousness (see Figure S3).

**Figure 5.**
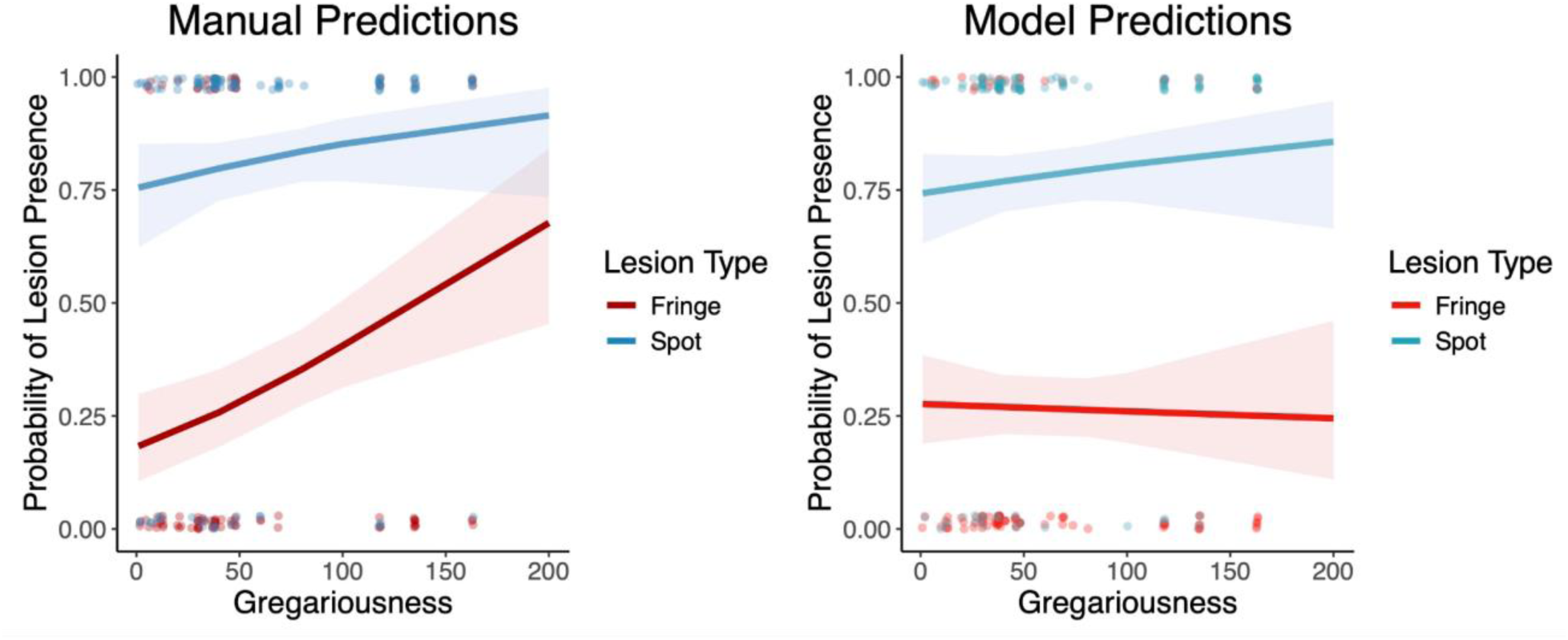
Individual-level analyses based on automated lesion detection should be cautious of data quality. The effect of gregariousness on the presence of lesions on individuals based on the manual determination of lesion presence (left) and the model determination of lesion presence (right). The model predictions are unable to identify the effect between gregariousness and lesion presence due to correlations between the independent variable (gregariousness) and data quality.

## 4 Discussion

We developed an efficient small object detection model for detecting lesions on dolphin skin to support rapid and effective health monitoring in delphinid species. We found that a high object-to-background ratio (via images cropped to contain the dolphin only, with minimal background) and partitioning of detection by lesion type were key to high model performance. Since manual image cropping to maximize object coverage is labor intensive, we developed an object detection model that automated the cropping of dolphin body and dorsal fin from images taken in the field.

We found that this automated cropping procedure is effective at increasing the object-to-background ratio and thus significantly increased our model performance. When conducting out-of-sample testing of our best detection model, we found that our model produced prevalence estimates that are consistent with manual estimates but is more sensitive to data quality on the individual-level association between gregariousness and lesion prevalence.

We found that data quality is central to strong model performance, particularly for spot lesions, as they are difficult to classify in photos or can be confused with water droplets and shadows. Thus, improving image quality and image replication to determine lesion presence improved model accuracy. Data quality may have less of an impact on the model’s ability to accurately predict fringe ring lesions since these rings are easier to identify even for human researchers (Toms et al., 2020). Indeed, the model’s precision and accuracy scores were higher when predicting the presence of fringe rings compared to spots. However, our sample of dolphins with fringe ring lesions in these high-quality images was small, so we were unable to reliably estimate the true effect of photo quality for this lesion type. The impact of data quality is apparent in our analysis of gregariousness. Our model tended to be less accurate at predicting lesions on individuals with larger gregariousness scores, which were negatively associated with the number of photographs of an individual provided to the model. Since our model is currently only equipped to detect lesions on images with one dolphin present, this effect is likely due to researchers having fewer opportunities to obtain multiple photographs of the same individual when group sizes are large, which subsequently impacts the model’s accuracy. We therefore suggest that users of this model prioritize high-quality images and endeavor to have at least three photographs of each dolphin to detect accurate lesion presence. We further conclude that our model may be reasonably used for population-level ecological questions but must be deployed with caution on individual-level ecological questions in which the data quality may covary with the variable of interest (e.g., social parameters).

In this work, we estimate a ∼70-85% prevalence of spot lesions and a ∼20-30% prevalence of fringe ring lesions on the dolphins that occupy the PCDP study area. Comparatively, estimates of lesions from other study sites can range from values as low as 14-39% prevalence in Redfish Bay, Texas during summer months (Guinn et al., 2024) to as high as 90.5-100% in NE Scotland (Wilson et al., 2000). Among sample sites along the East Coast of the United States, prevalence seems to fall close to 50% (Guinn et al., 2024; Hart et al., 2012; Taylor et al., 2021), suggesting that the prevalence of spot lesions (and thus lesions in general) may be elevated for animals in the Potomac-Chesapeake. This could be because many animals that use the Chesapeake are part of migratory populations (Hayes et al., 2023) that are exposed to different habitats, temperatures, and salinities, which influence the presence of skin lesions in dolphins (Duignan et al., 2020; Stylos et al., 2022; Wilson et al., 1999), and the Chesapeake Bay itself has distinct geographic salinity divisions and annual temperature fluctuations (Baird & Ulanowicz, 1989). Areas such as our study site in the Potomac River also contain pollution from agricultural runoff, wastewater, and other sources, which can contribute to weakening the dolphins’ immune systems, making them vulnerable to more common viral diseases as well as potential adverse health conditions from toxin accumulation (Bennett et al., 2001; Koch et al., 2018; Lahvis et al., 1995; Page-Karjian et al., 2020; Pierce et al., 2008). Other stressors that could also weaken dolphin immune systems include high levels of human activity — which can cause noise disturbance, injury, and prey depletion — and freshwater input from storms that reduce salinity and increase lesion prevalence (Collier et al., 2022; Deming et al., 2020; Duignan et al., 2020; Fazioli & Mintzer, 2020).

Our estimates of lesion prevalence begin to shed light on the potential health of the bottlenose dolphins in this region. While the etiology of spot lesions remains unknown, there is growing evidence for fringe ring lesions as indicators of infectious diseases (Geraci et al., 1979; Hart et al., 2012; Toms et al., 2020). Indeed, we were able to determine a positive effect of group size on the presence of fringe ring lesions on individuals but were unable to do so with spot lesions. This suggests that more gregarious individuals could be at higher risk for contracting infectious skin diseases (as indicated by fringe ring lesions), as previously found in other bottlenose dolphins (Powell et al., 2020). Gregariousness can also vary across different populations of dolphins that range in size (Read et al., 2013). Since both smaller estuarine and larger migratory populations visit our study site seasonally (Hayes et al., 2023), individuals from these larger populations likely make up the higher gregariousness values in our study. Individuals from these populations may be at greater risk of infection due to their gregariousness and migratory behaviors, exposing them to greater environmental fluctuations. These results contrast our alternative hypothesis that visible infections may cause avoidance by conspecifics or sickness behaviors by individuals, which are seen across many species (Stockmaier et al., 2023). While our results do not support this hypothesis, our measure of gregariousness (average survey size each individual was sighted in) is broad, and this analysis should be expanded on with other social metrics (e.g., average number of close or physical associates) to provide stronger evidence.

Our model performs well in addressing ecological questions (under conditions of high data quality), suggesting that it can serve both as a powerful tool in its own right and an excellent starting point for any further additions or optimizations. Future work could build on our model to recognize more lesion types. Efforts to predict other lesions of interest such as tattoo skin disease lesions (Geraci et al., 1979; Maldini et al., 2010; M. F. Van Bressem et al., 1999) or other visible signs of disease and poor health such as lobomycosis (Reif et al., 2009; Toms et al., 2020) would allow for researchers to get an accurate estimate of disease prevalence in their populations in a short amount of time compared to manual methods. From the 1987 and 2013 dolphin morbillivirus outbreaks that depleted more than half of some Atlantic bottlenose dolphin populations (Lipscomb et al., 1994; Waring et al., 2016), we know that disease outbreaks in dolphins can have devastating outcomes. Real-time monitoring of skin lesions associated with infectious diseases could be used as an early warning sign of future deadly outbreaks and monitoring individual behaviors alongside skin lesions could be used to help model disease transmission. Object detection models such as ours can improve the efficiency with which health assessments are conducted for conditions apparent from external observation. Disease monitoring in wildlife is of critical importance, especially in marine mammals, whose health has been declining over recent decades (Gulland & Hall, 2007) and is projected to worsen further due to climate change (Gulland et al., 2022; Sanderson & Alexander, 2020). Skin diseases in wildlife have also been observed in terrestrial wildlife such as red foxes (Carricondo-Sanchez et al., 2017) and wolves (Oleaga et al., 2011), suggesting that similar models could improve the efficiency of health monitoring for a wide variety of vulnerable taxa.

## Author Contributions

Conceptualization: CJM, MAC, SB

Data Collection: MAC, A-MJ, EK, MMW, JM

Analysis Tool Contribution: CJM, MAC, SB

Analysis: CJM, MAC, SB

Supervision: MAC, SB

Funding Acquisition: MAC, A-MJ, EMP, JM, SB

Writing—original draft: CJM

Writing—review & editing: MAC, A-MJ, EMP, MMW, JM, SB

## Statement on Inclusion

Our study was carried out near Washington, DC, United States and thus brings together authors from both academic and federal institutions. Data collection and analysis was carried out by graduate student researchers, and undergraduate research assistants who were brought on from the local DC area and offered paid stipends to foster an inclusive and accessible environment. Our final model is provided free to use for researchers of any institution world-wide.

## Acknowledgements and Funding

We thank Roboflow for providing access to their platform for model creation and assessment, and are grateful to Mohamed Traore for his assistance with the platform throughout this project. We thank Denise Greig and the Marine Mammal Health and Stranding Response Program for their feedback on this work. We would also like to thank Milan Dolezal, Amelia Smith, and Katherine Dammer for their assistance with lesion coding on our dataset. All data from the PCDP was collected under NMFS Permit nos 19403 and 23782. This work was supported by the Morris Animal Foundation Award #D22ZO-059, and funding from Georgetown University. Data collection for this work by the PCDP was also supported by the Potomac River Keepers, the Rogers Family Foundation, the Campbell Foundation, the Scheidel Foundation, Georgetown University Earth Commons, Green-Rosenblum Family Foundation, the Wildlife Conservation Society, the National Geographic Society Grant WW-022ER-17, Waldorf Toyota, and individual donors.

## Data Availability Statement

All data and code files have been uploaded to https://doi.org/10.5281/zenodo.12735326. If accepted, we plan to host these data and code files in a publicly accessible repository.

