## Supplement for "Automated Skin Lesion Detection and Prevalence Estimation in Tamanend’s Bottlenose Dolphins"

### Transfer Learning

Transfer learning is the process of initializing the network's weights with the weights from a previous model. This can be done using a standardized model like COCO or ImgNet, where the weights for layers closer to the output layer will need to be trained but basic layers such as edge detection are already well established. It can also be done using previous iterations of the model being trained, or with a specialized prototype model designed to help set the weights to extract useful information in a controlled environment. When we used transfer learning in our project, we used this last method of transfer learning. Verification after the project was completed showed that the transfer learning did not seem to significantly impact training results, but was kept for consistency across models.

### Preprocessing and Augmentation

When working with large uncropped images, we applied a preprocessing step known as tiling. We tiled all uncropped images with a 6x9 grid, such that all tiles were squares with 608 pixels per side (Figure 2). These tiles were then considered the source images used for all future steps of the model's design and training process. For cropped images in the one-step process, no tiling was performed. For both lesion detectors specialized to detect on the body crops of the two-step model, tiling was done along a 1x4 grid, as the mean aspect ratio of these crops was roughly four times as wide as it was tall.

We then created five augmented images from each source image to 1) create “new” training images from existing training images to increase the amount of data available for the model to learn from and 2) allow the model to better prepare for elements that are not present in the training data. Augmentation methods included a random crop of 0-20% to simulate different zooms, random rotation of 0-15% in any direction and random shear of 0-5% in any direction to simulate photographing the dolphins from different angles. Augmentations also included random blur up to 0.5 pixels and random noise in up to 1% of all pixels to simulate blurriness or splashes interfering with the image. Finally, mosaic augmentation was used as well for the augmented images, which pieces together parts of different training images to form new training images.

**Table S1.** Broad overview of data used to train lesion detection models.

| Year | Number of Surveys | Number of Photos |
|---|---|---|
| 2015 | 12 | 230 |
| 2016 | 12 | 219 |
| 2017 | 18 | 263 |
| 2018 | 14 | 191 |

**Table S2.** Years from which data was sourced when constructing each of our models. Note that photos in the one-step cropped, fringes only, and spots only models may be multiple crops capturing different dolphins in the same photo, explaining some of the increase in number of photos over the uncropped model.

| Dataset | Photos - 2015 | Photos - 2016 | Photos - 2017 | Photos - 2018 | Total |
|---|---|---|---|---|---|
| One-Step Uncropped | 193 | 154 | 58 | 0 | 405 |
| One-Step Cropped | 247 | 228 | 269 | 213 | 957 |
| One-Step Fringes Only | 228 | 184 | 218 | 183 | 813 |
| One-Step Spots Only | 241 | 183 | 226 | 188 | 838 |
| Two-Step Dorsal Fin Fringes Only | 119 | 132 | 153 | 106 | 510 |
| Two-Step Dorsal Fin Spots Only | 118 | 132 | 153 | 106 | 509 |
| Two-Step Body Fringes Only | 114 | 131 | 134 | 102 | 481 |
| Two-Step Body Spots Only | 108 | 131 | 130 | 100 | 469 |
| Test Dataset | 156 | 88 | 122 | 0 | 366 |

**Table S3.** Image quality scoring for photographs in the PCDP. Grading criteria (focus, visibility, contrast, and angle) from Urian et al., 2015. Table from Dolezal et al., 2023.

**Grade 0**

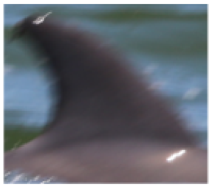

**Poor quality.** The dorsal fin is: 1) out of focus and extremely blurry or pixelated; 2) not visible or 10% or less of the fin is visible; 3) extremely under or overexposed; and/or 4) at an extreme acute or obtuse angle to the camera.

**Grade 1**

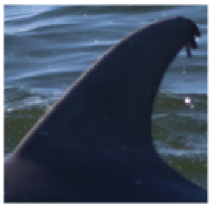

**Moderate quality.** The dorsal fin is: 1) out of focus and blurry or pixelated; 2) less than 90% visible; 3) under or overexposed; and/or 4) at an acute or obtuse angle to the camera.

**Grade 2**

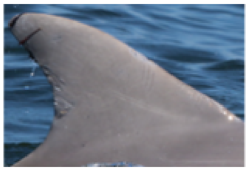

**Good quality.** The dorsal fin is: 1) very slightly out of focus or pixelated; 2) 90% or more visible; 3) very slightly under or over exposed; and/or 4) at a very slight acute or obtuse angle to the camera.

**Grade 3**

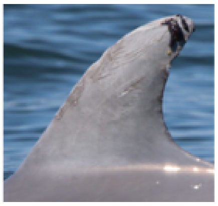

**Excellent quality.** The dorsal fin is: 1) in sharp focus; 2) fully visible; 3) well exposed; and 4) at a right angle to the camera.

**Table S4.** Roboflow output for models which located dolphin's body and dorsal fin.

| Replicate | mAP | Precision | Recall |
|---|---|---|---|
| v1 | 94.1% | 95.2% | 85.3% |
| v2 | 97.8% | 96.0% | 94.1% |
| v3 | 96.8% | 94.6% | 93.3% |
| v4 | 96.3% | 96.9% | 91.6% |
| v5 | 98.2% | 98.5% | 94.1% |
| v6 | 96.6% | 96.7% | 92.7% |
| v7 | 97.1% | 96.9% | 94.3% |
| v8 | 97.7% | 95.4% | 94.6% |
| v9 | 97.6% | 95.4% | 95.7% |
| v10 | 97.5% | 93.8% | 94.9% |

**Table S5.** Result of TukeyHSD test performed comparing model performance statistics.

| Pair | Diff | p adj |
|---|---|---|
| Uncropped / Cropped | -42.00 | 0.0000000 |
| Uncropped / Cropped Spots | -30.85 | 0.0000000 |
| Uncropped / Cropped Fringes | -64.31 | 0.0000000 |
| Cropped / Cropped Spots | 11.15 | 0.0000000 |
| Cropped / Cropped Fringes | -22.31 | 0.0000000 |
| Cropped Spots / Cropped Fringes | -33.46 | 0.0000000 |
| Cropped Spots / Body Spots | -33.76 | 0.0000000 |
| Cropped Spots / Body Fringes | -42.56 | 0.0000000 |
| Cropped Spots / Dorsal Fin Spots | -30.15 | 0.0000000 |
| Cropped Spots / Dorsal Fin Fringes | -39.61 | 0.0000000 |
| Cropped Fringes / Body Fringes | -9.10 | 0.0000016 |
| Cropped Fringes / Body Spots | -0.30 | 0.9999993 |
| Cropped Fringes / Dorsal Fin Fringes | -6.15 | 0.0027080 |
| Body Spots / Body Fringes | -8.80 | 0.0000037 |
| Body Spots / Dorsal Fin Spots | 3.61 | 0.2577219 |
| Body Spots / Dorsal Fin Fringes | -5.85 | 0.0052203 |
| Body Fringes / Dorsal Fin Fringes | 2.95 | 0.5152414 |
| Body Fringes / Dorsal Fin Spots | 12.41 | 0.0000000 |
| Dorsal Fin Spots / Dorsal Fin Fringes | -9.46 | 0.0000006 |

**Table S6.** Confusion matrices for model predictions separated by lesion class and number of photos per dolphin. All numbers for this set of confusion matrices represent the number of dolphins not the number of images.

| Spots, 1 Photo | **Actual Positive** | **Actual Negative** |
|---|---|---|
| **Model Positive** | TP: 78 | FP: 12 |
| **Model Negative** | FN: 28 | TN: 14 |

| Spots, 2 Photos | **Actual Positive** | **Actual Negative** |
|---|---|---|
| **Model Positive** | TP: 49 | FP: 6 |
| **Model Negative** | FN: 4 | TN: 1 |

| Spots, 3+ Photos | **Actual Positive** | **Actual Negative** |
|---|---|---|
| **Model Positive** | TP: 24 | FP: 2 |
| **Model Negative** | FN: 2 | TN: 0 |

| Fringes, 1 Photo | **Actual Positive** | **Actual Negative** |
|---|---|---|
| **Model Positive** | TP: 19 | FP: 8 |
| **Model Negative** | FN: 25 | TN: 80 |

| Fringes, 2 Photos | **Actual Positive** | **Actual Negative** |
|---|---|---|
| **Model Positive** | TP: 14 | FP: 7 |
| **Model Negative** | FN: 7 | TN: 32 |

| Fringes, 3+ Photos | **Actual Positive** | **Actual Negative** |
|---|---|---|
| **Model Positive** | TP: 9 | FP: 2 |
| **Model Negative** | FN: 4 | TN: 13 |

**Table S7.** Statistics for model performance based on the number of photos per dolphin.

| Subdivision | Recall | Precision | Accuracy | F-Score |
|---|---|---|---|---|
| Spots, 1 Photo | 73.6% | 86.7% | 69.7% | 79.6% |
| Spots, 2 Photos | 92.5% | 89.1% | 83.3% | 90.7% |
| Spots, 3+ Photos | 92.3% | 92.3% | 85.7% | 92.3% |
| Fringes, 1 Photo | 43.2% | 70.4% | 75.0% | 53.5% |
| Fringes, 2 Photos | 66.7% | 66.7% | 76.7% | 66.7% |
| Fringes, 3+ Photos | 69.2% | 81.8% | 78.6% | 75.0% |

**Table S8.** Confusion matrices for model predictions separated by lesion class and image quality rating of photo. All numbers for this set of confusion matrices represent images not dolphins.

| Spots, 2 Stars | **Actual Positive** | **Actual Negative** |
|---|---|---|
| **Model Positive** | TP: 179 | FP: 63 |
| **Model Negative** | FN: 66 | TN: 41 |

| Spots, 3 Stars | **Actual Positive** | **Actual Negative** |
|---|---|---|
| **Model Positive** | TP: 17 | FP: 0 |
| **Model Negative** | FN: 0 | TN: 0 |

| Fringes, 2 Stars | **Actual Positive** | **Actual Negative** |
|---|---|---|
| **Model Positive** | TP: 42 | FP: 26 |
| **Model Negative** | FN: 45 | TN: 236 |

| Fringes, 3 Stars | **Actual Positive** | **Actual Negative** |
|---|---|---|
| **Model Positive** | TP: 4 | FP: 0 |
| **Model Negative** | FN: 5 | TN: 8 |

**Table S9.** Statistics for model performance based on photo quality

| Subdivision | Recall | Precision | Accuracy | F-Score |
|---|---|---|---|---|
| Spots, 2 Stars | 73.1% | 74.0% | 63.0% | 73.5% |
| Spots, 3 Stars | 100% | 100% | 100% | 100% |
| Fringes, 2 Stars | 48.3% | 61.8% | 79.7% | 54.2% |
| Fringes, 3 Stars | 44.4% | 100% | 70.6% | 61.5% |

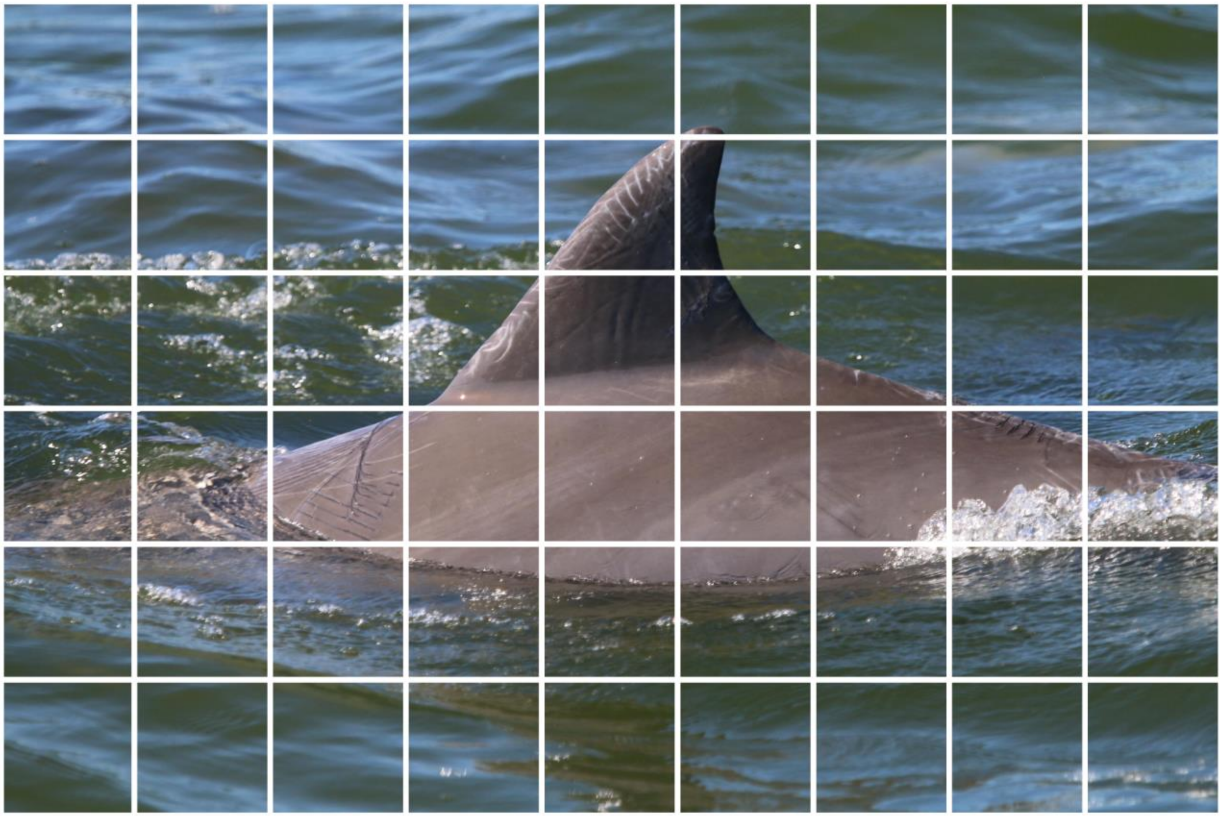

**Figure S1.** Example of a photograph tiled on a 6x9 grid. This is one of our 5472x3648 px photographs taken from the field, tiled into 608x608 px squares. Each of the 54 resulting squares was treated as a source image for the development of the model. In any square containing a lesion, the area of the lesion would take up a larger proportion of the square than it would with the full image, making it easier to detect.

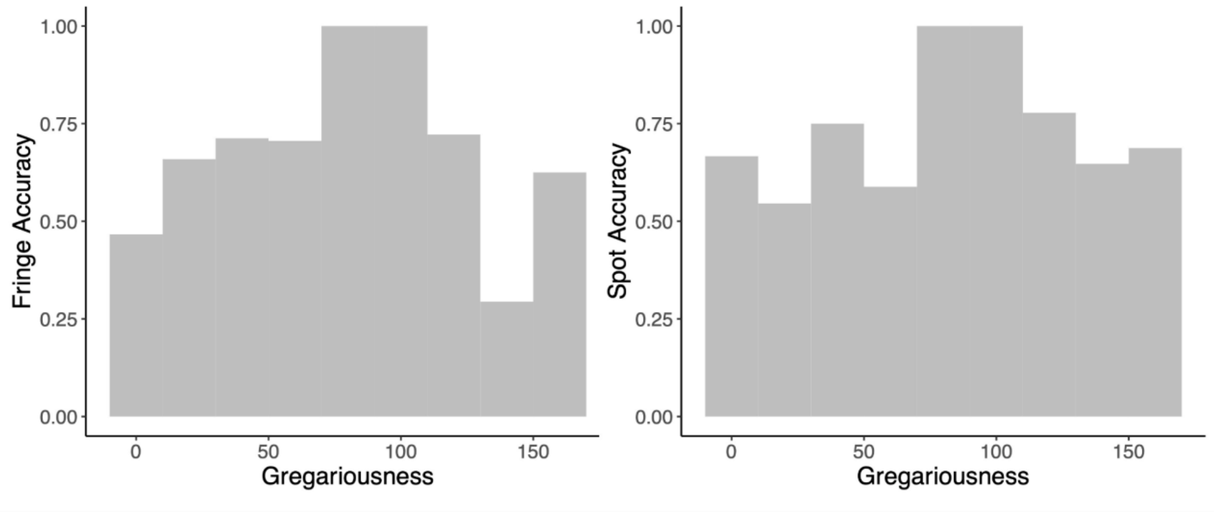


**Figure S2**: The accuracy of the model estimates for lesions presence on individuals across sociality levels for fringe (left) and spot (right).

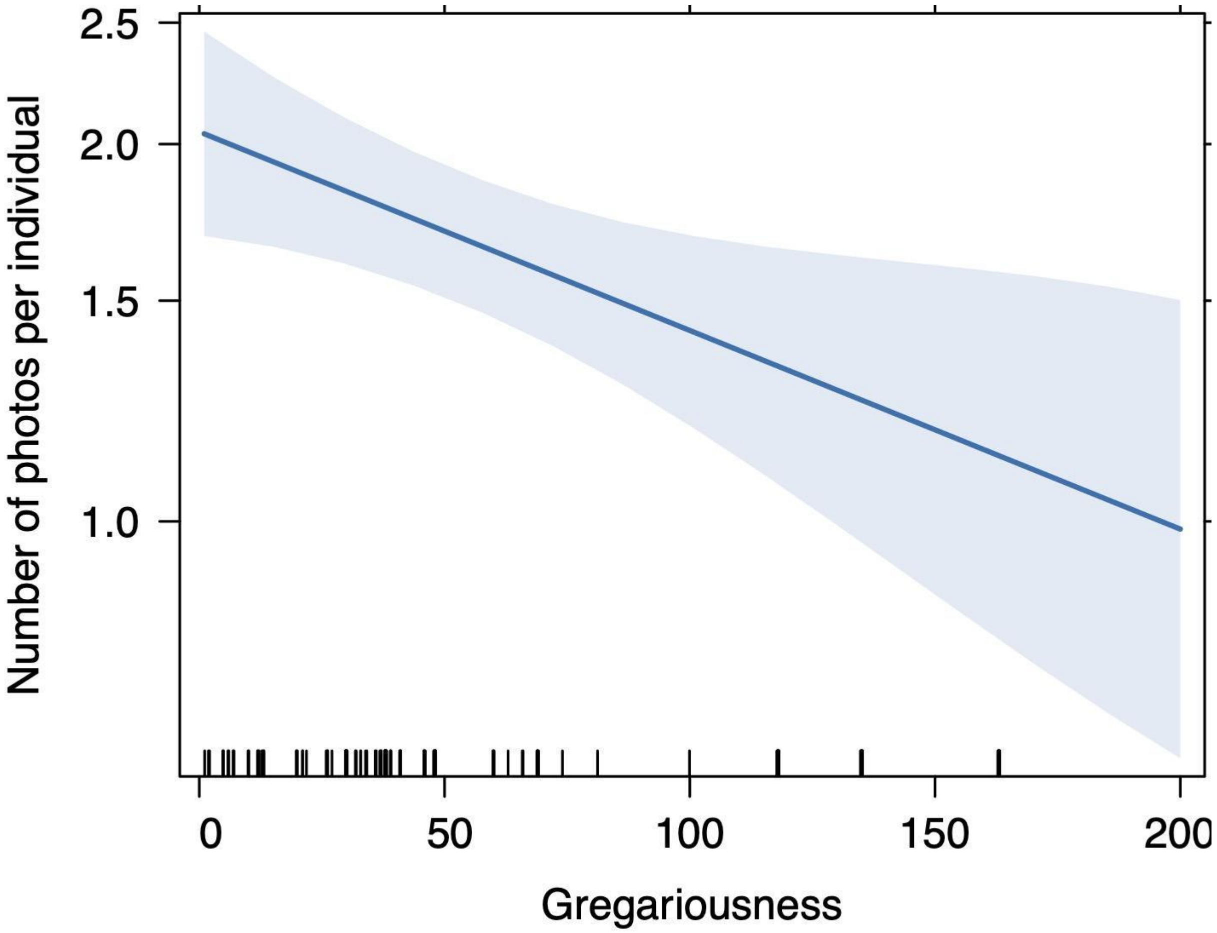


**Figure S3**: The effect of sociality on the number of photos per individual ran through the model. We see a significant negative effect of sociality ($z = -2.609$ , $p = 0.00909$) suggesting that individuals seen in larger survey sizes have less individual photographs of them available to be run through our model.
